# 4-Pi Stimulated Raman Scattering for Label-free Super-resolution Chemical Imaging

**DOI:** 10.1101/2025.08.28.672954

**Authors:** Jonathan I. Kim, Zachary Ellsworth, Erin L. Dunnington, Nidhi R. Mehta, Chisa Zensho, Dan Fu

**Affiliations:** Department of Chemistry, University of Washington, Seattle, WA 98195, United States

## Abstract

Super-resolution fluorescence microscopy has transformed the study of biological structures and functions beyond the diffraction limit. Unlike fluorescence methods, label-free chemical imaging, mostly based on Raman and infrared spectroscopy, provides intrinsic molecular contrast, enabling the study of biomolecules, nanostructures, drug molecules, and metabolites that cannot be easily tagged. However, while a wide range of fluorescence-based super-resolution techniques are well-established, extending super-resolution to label-free chemical imaging has remained challenging due to low signal levels and limited resolution improvement. Super-resolution stimulated Raman scattering microscopy (SRS) is most promising due to its high sensitivity and imaging speed. Similar to fluorescence, existing SRS super-resolution approaches are mostly based on either photoswitching/saturation of molecular labels or sample expansion, which suffers from poor sensitivity due to limitations in labeling density or signal dilution, respectively. Moreover, axial resolution is typically much worse than lateral resolution, yet most super-resolution SRS techniques focused on improving lateral resolution. In this work, we combine stimulated Raman scattering (SRS) with 4Pi-interferometry to significantly improve the axial resolution by nearly 7-fold. We report on the characterization of improvements in imaging sensitivity and axial resolution using 80 nm polystyrene beads. Harnessing the improved axial resolution, we demonstrate super-resolution 4Pi-SRS imaging in resolving small lipid droplet structures in mammalian cells and lipid membranes in *E. coli* cells. Because 4Pi-SRS uses interferometry to improve axial resolution, it is completely orthogonal to all previous super-resolution SRS techniques, including visible excitation, photoswitching, sample expansion, and computational approaches, thus it is straightforward to combine them to achieve much higher resolution chemical imaging than currently possible.

## INTRODUCTION

The diffraction limit has long constrained the ability to resolve fine biological structures, causing many microscopic observations to remain blurred in conventional microscopy.^1^ The development of super-resolution fluorescence microscopy broke the diffraction barrier by either re-engineering the excitation point spread function or exploiting fluorophore ON/OFF switching to distinguish emitters within the diffraction-limited excitation volume.^2,3^ These advances have yielded transformative insights into the nanoscale organizations and functions of proteins and organelles, reshaping our understanding of cellular architecture and dynamics. While genetically encoded or chemically engineered fluorophores have made it possible to visualize many biomolecules, many important molecular targets, such as small drug molecules and metabolites, remain inaccessible. Fluorescent labeling is also subject to other limitations including color barriers, photobleaching, and phototoxicity. Consequently, large areas of cellular biochemistry remain invisible to current super-resolution fluorescent techniques. This unmet need highlights the importance of developing label-free chemical imaging approaches that can directly probe molecular composition and dynamics without relying on exogenous tags, thereby extending the super-resolution principles to a broader range of biological structures and functions.

Stimulated Raman scattering (SRS) microscopy serves as a powerful platform for super-resolution label-free chemical imaging.^4,5^ Utilizing synchronized pump and Stokes lasers to coherently drive a vibrational transition, SRS enables high resolution imaging at speeds orders of magnitude faster than spontaneous Raman scattering.^6,7^ SRS has been transformative in mapping cell metabolism and metabolites, quantifying drug uptake in live cells, imaging neurotransmitters, stimulated Raman histology, and super-multiplexed vibrational imaging with >20 colors, among many other applications.^7–13^ Despite the breadth of applications, SRS would benefit from resolution gains both laterally and axially.

Resolution enhancement in SRS has been pursued through strategies inspired by super-resolution fluorescence microscopy. However, unlike fluorescence with single molecular sensitivity, SRS sensitivity is at best in the micromolar range, far from single molecule. It is challenging if not impossible to rely on localization based super-resolution. Instead, shrinking the excitation volume using stimulated emission depletion or photoswitching becomes the favored choice. The use of “donut-shaped” illumination similar to stimulated emission depletion was first demonstrated in nonbiological samples.^14^ However, depletion of SRS required extremely high power that is incompatible with biological imaging. The problem was addressed through depletion of stimulated Raman-excited fluorescence for fluorescent labeled samples.^15^ Coupling the depletion approach with reversibly photoswitching Raman probes allows detection of SRS from those probes at a much small volume confined by the gaussian activation beam and donut depletion beam. The power required for depletion is much lower and biological imaging at resolution < 100 nm was shown to be possible.^16,17^ However, these approaches all require special labels. Other strategies based on nonlinear effects —such as saturation,^18–21^ higher-order susceptibilities,^22^ and structured illumination ^23–25^—have also been shown to provide up to ∼2-fold lateral resolution improvements. Saturation and higher-order approaches trade off photodamage for resolution, leading to biocompatibility issues. The more straightforward way to improve resolution is through visible excitation. Although diffraction limited, frequency-doubling of near-infrared SRS excitation into the visible directly improves the resolution by 2-fold.^26^ Further extension through Fourier reweighting achieved ∼130 nm and ∼85 nm lateral resolution, respectively.^27^ Computational approach based on deconvolution has recently been applied to sparse features, resolving lipid droplets at ∼60 nm,^28^ though it is difficult to evaluate the quantitiveness of the results and there is potential for artifacts especially with weak signals. Physical expansion using swellable hydrogels offers another route to improve effective resolutions below 100 nm.^29,30^ Despite these advances, most of the aforementioned methods focus on improving lateral resolution, yet like most light microscopy modalities, SRS suffer from poorer axial than lateral resolutions, often 3-5 times worse.^31^ The diffraction limit on the axial direction is much more difficult to overcome than the lateral direction.

It is possible to use a 3D stimulated emission depletion approach to achieve super-resolution in both the lateral and axial dimensions. However, a fundamental constraint of labeling based super-resolution SRS imaging is the limited sensitivity. Unlike super-resolution fluorescence microscopy that detects individual fluorescent emitters, SRS sensitivity is typically at millimolar or high micromolar level even with strong Raman dyes. SRS signal is linearly proportional to concentration and excitation volume. Shrinking the excitation volume is bound to suffer from signal loss. The labeling density thus poses a limit to the achievable resolution. Similarly, high resolution SRS based on sample expansion further dilutes already weak SRS signals, making it difficult to resolve nanostructures.^29,30^ Visible SRS does not suffer from those problems, however, it is limited by phototoxicity at shorter wavelengths, which comes at the cost of sensitivity.^26,27^

4Pi-fluorescence microscopy is a well-established method, shown to both improve the axial resolution and sensitivity for fluorescence.^32–34^ A doubling of the collection efficiency improves the sensitivity ∼1.4-fold, and axial interference improves the axial resolution ∼3-7-fold. While 4Pi-interferometry has been combined with spontaneous Raman imaging,^35^ its application was limited to thin homogeneous thin layer substrates and no imaging was performed. Here, we combine SRS with 4Pi-interferometry, achieving ∼7-fold axial resolution improvements (from ∼1000 nm to ∼142 nm) and 2-4x sensitivity enhancement. We characterize the axial resolution improvements of the 4Pi-SRS microscope by performing three-dimensional imaging of 80 nm polystyrene beads, showing a super-resolved axial profile of 155 nm full-width half-maximum (FWHM). To show the benefits of the 4Pi-SRS microscope in biological imaging, we demonstrate substantial contrast improvement in lipid droplet imaging, with an axial profile of 160 nm FWHM. Additional deconvolution allows for side lobe suppression for super-resolved lipid droplet structures. Finally, we perform three-dimensional 4Pi-SRS deconvolution imaging of *E. coli* membrane and fatty acid bioproduct. The unique signal enhancement and axial resolution improvement of 4Pi-SRS is complementary to other resolution enhancement techniques; thus, it would be straightforward to combine 4Pi-SRS with visible excitation, sample expansion, or photoswitching to further push the resolution limit to <100 nm in all three dimensions. Such capabilities would open up new avenues of super-resolution imaging of biological structure and function.

## Results

### 4Pi-SRS Microscope Design

To realize the 4Pi-SRS microscope system, we integrated a conventional SRS setup with a 4Pi interference cavity constructed by two opposing high numerical aperture (NA) objectives, as shown in Figure 1a. The layout of the 4Pi cavity is similar to that implemented in two-photon 4Pi-fluorescence microscopy.^34^ The main difference is that SRS employs two excitation lasers, pump and Stokes. The focus and interference of both excitations have to overlap exactly in the axial direction for optimal SRS excitation. For SRS imaging, the Stokes beam centered at 1030 nm is modulated at 17.5 MHz with an electro-optic modulator (EOM), enabling detection of the stimulated Raman loss in the pump beam at 793 nm. The two beams are spatially overlapped using a dichroic mirror (DM) before being sent into the 4Pi cavity. The temporal delay between pump and Stokes is controlled by a motorized stage on the pump beam path for acquiring hyperspectral SRS images via spectral focusing.^36^

**Figure 1.**
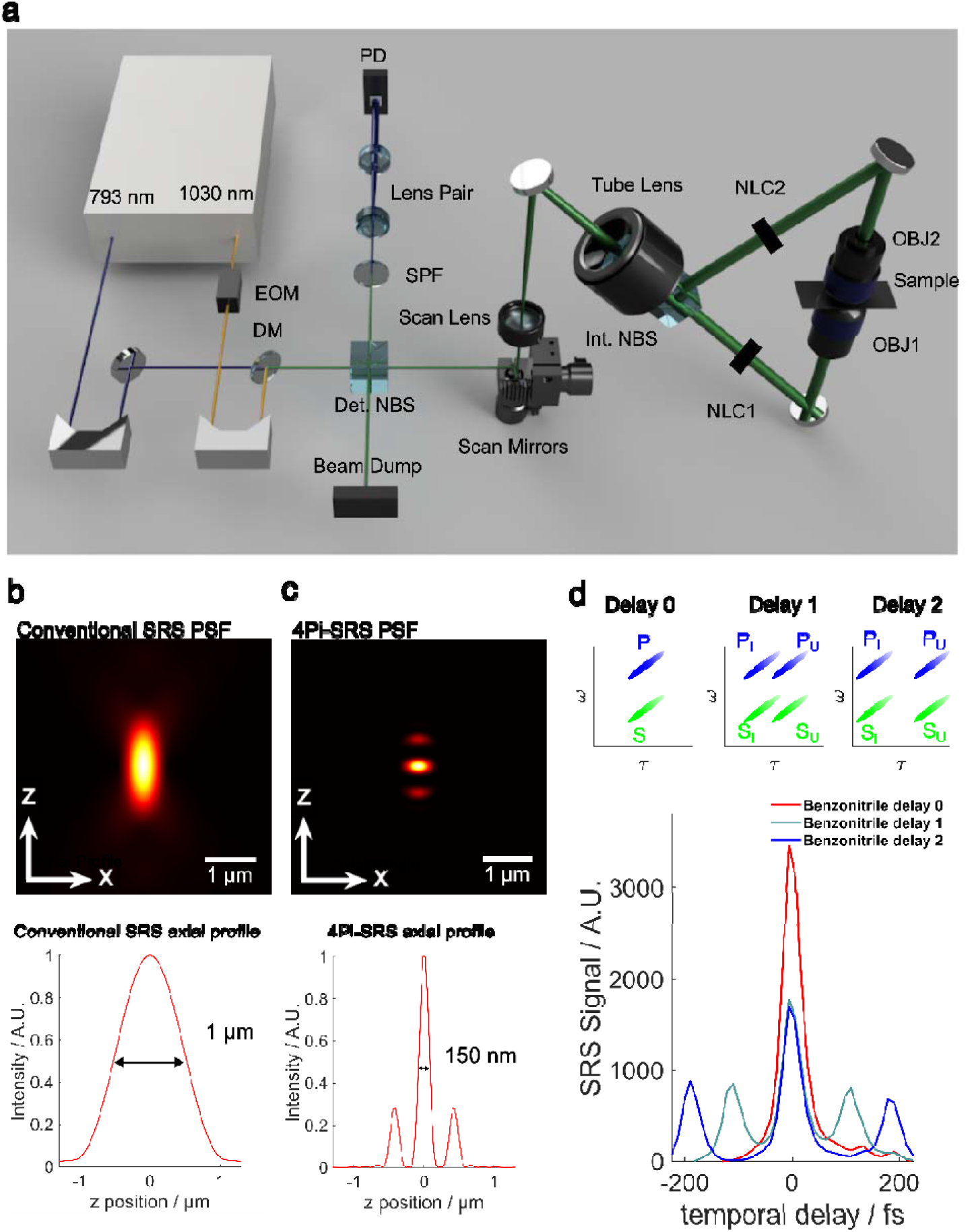
(a) 4Pi-SRS microscope diagram. EOM = electro-optic modulator; 50/50 NBS = 50/50 non-polarizing beam-splitter; DM = dichroic mirror; SPF = short-pass filter; PD = photodiode detector; QWC = quartz-wedge compensator; OBJ = objective, NLC = nematic liquid crystal. (b) Simulated 3D conventional SRS PSF (top) and its axial profile (bottom), and (c) simulated 4Pi-SRS PSF (top) and its axial profiles (bottom). (d) SRS spectra of benzonitrile ∼3070 cm□^1^ Raman transition of aromatic CH stretching. Delay 1 indicates a slight optical path length mismatch between he upper and lower arms, while Delay 2 corresponds to a larger mismatch. P_l_ = lower arm pump, P_u_ = upper arm pump, S_l_ = lower arm Stokes, S_u_ = upper arm Stokes.

The 4Pi configuration employs two 50/50 non-polarizing beam splitters (NBS). The first denotes the detection NBS as it directs the transmitted pump beam after interference to the SRS detector. The second NBS, the interference NBS, equally splits the pump and Stokes into two arms — an upper and lower arm, forming the 4Pi interference cavity. The focused pump and Stokes originating from the lower arm are collected through objective 2, while the pump and Stokes from the upper arm are collected through objective 1. Both propagate to the interference NBS, where the pump and Stokes constructively interfere and travel back to the detection NBS. At this point, half of the pump and Stokes are reflected by the detection NBS. The pump is then filtered and detected with a photodiode. To ensure that the two optical pathlengths from the NBS to the optical focus are perfectly matched for constructive interference of both pump and Stokes, two adjustments are necessary. One is the overall mechanical delay adjustment, which can be achieved by translating the top mirror above objective 2. The second adjustment is to fine tune the optical path length difference between pump and Stokes through voltage-controlled nematic liquid crystals (NLC1 and NLC2).

As a result of the interference cavity, both the counterpropagating pump and Stokes generate an axial interference fringe, shrinking the diffraction-limited excitation volume. To model this interference, we applied the diffraction theory model based on the works of Richards and Wolf to simulate the 3D point-spread function (PSF) found within Supplemental Note 1.^37,38^ The 3D PSF defines how a point source object is laterally and axially blurred, limiting resolution. The PSF of SRS is the multiplication of the PSFs for pump and Stokes. This simple model allows us to compare the resolution limits of conventional SRS and 4Pi-SRS. Using a 1.2 NA objective with pump and Stokes wavelength at 793 nm and 1030 nm, respectively, conventional SRS has a lateral resolution of ∼ 300 nm and axial resolution of ∼1 μm (Figure 1b). In contrast, the 4Pi-SRS 3D PSF exhibits axial interference fringes that result in a sharply defined central lobe, improving the effective axial resolution from ∼1000 nm to ∼150 nm, while maintaining the same lateral resolution (Figure 1c). The theoretical resolution improvement is ∼7-fold for the 4Pi-SRS microscope. The simulated side lobe intensity is approximately 30% of the central lobe, which can be removed computationally through deconvolution.^39–42^ A summarization of the deconvolution pipeline process is found within Supplemental Figure S1.

In addition to the axial resolution improvement, sensitivity enhancement is also expected because of the nonlinear SRS signal dependence on the peak intensity. We probed the ∼3070 cm^-1^ Raman transition of benzonitrile to quantify the sensitivity improvement. Since there are two counter propagating beam paths (upper to lower and lower to upper), four SRS processes can happen at the laser focus: I_p,l*_ I_s,l_, I_p,u*_ I_s,l_, I_p,l*_ I_s,u_, I_p,u*_ I_s,u_. Pulse pairs traveling in the same direction (I_p,l_* I_s,l_ and I_p,u_*I_s,u_) generate equal SRS signal at the same delay (defined as delay 0 in Figure 1d) in the spectral focusing scheme. However, when the upper arm path length is longer than the lower arm, to probe the same Raman transition, I_p,l_*I_s,u_ will generate SRS signal at positive delay (increasing pump travel distance), while I_p,u_*I_s,l_ will generate SRS signal at the negative delay. We define these two signals as the cross-arm SRS signals. As a result, three SRS peaks will appear as the pump and Stokes delay is scanned. The two sidelobes have half the intensity of the central peak because there is only one pair of interaction as opposed to two pairs. The separation of these three transitions reflects the mismatches in the optical path lengths between the two arms (shown as delay 1 and delay 2). When path lengths are perfectly matched, the three transitions merge into one and lead to a two-fold increase in absolute SRS signal. The limit of detection for 4Pi-SRS measurements with dimethyl sulfoxide (DMSO) sample can be found in Supplemental Figure S2.

### Characterization of 4Pi-SRS Resolution

To evaluate the resolution improvement of 4Pi-SRS over conventional SRS, we experimentally quantified its axial resolution by imaging polystyrene (PS) beads of varying diameters at the ∼3050 cm^-1^ Raman transition. Bead sizes were chosen to compare object-limited versus resolution-limited profiles for the 4Pi-SRS PSF.

To assess the 4Pi-SRS PSF, we first imaged 80 nm PS beads, which are smaller than the diffraction-limited axial resolution. The axial and lateral intensity of a representative bead (Figure 2a-c) were fit using three-peak Gaussian and single-peak Gaussian fitting respectively, yielding an axial FWHM of 155 nm and an x-lateral FWHM of 316 nm. An average of eight 80 nm beads were taken, yielding an average axial FWHM of 162 ± 13 nm, and an x-lateral FWHM of 323 ± 34 nm. These 80 nm beads were resolved with an average signal-to-noise ratio (SNR) of 7 before denoising, the smallest beads resolved with near-infrared excitation SRS to our knowledge. In comparison, conventional SRS of these beads do not have sufficient SNR for FWHM characterization. Previously sub-100nm beads are only detectable with visible SRS.^26^ We note that the experimentally observed side lobes reached ∼40% of the central lobe intensity, slightly higher than the ∼30% predicted by simulations. This is likely due to residual spherical or chromatic aberration of the imaging system. The 80 nm bead profile can be used as an estimation of the 4Pi-SRS PSF, given that the bead size is smaller than the axial and lateral resolution limits of the microscope. The 80 nm bead measurements were highly reproducible in their axial interference pattern as shown in Supplemental Figure S3. We estimated the resolution of the 4Pi-SRS system to be ∼142 nm based on the measured bead size, which is a convolution of the PSF of 80 nm bead with its true size.

**Figure 2.**
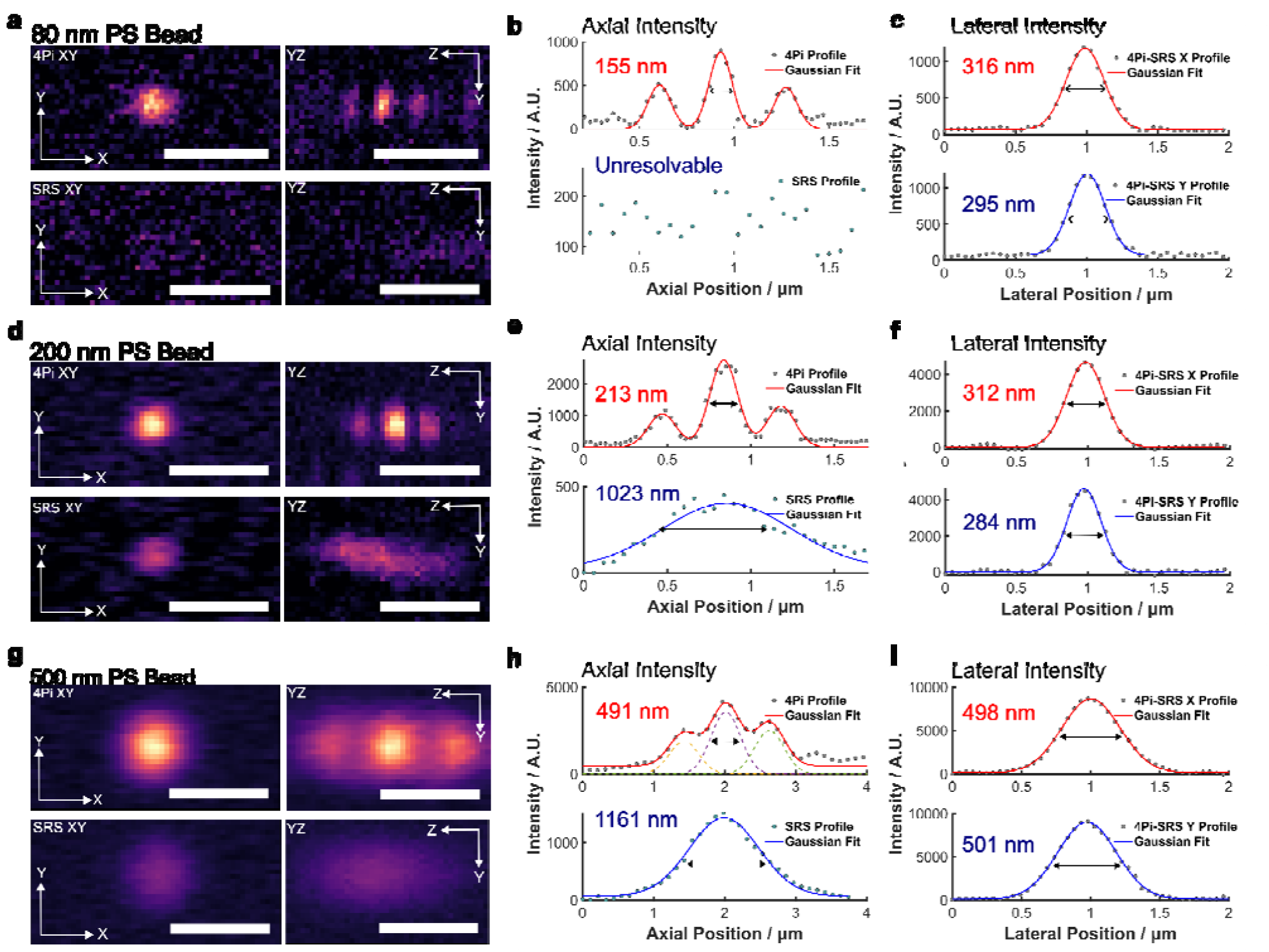
(a, d, g) Comparison of 4Pi and conventional SRS imaging of polystyrene beads. (b, e, h) Axial profiles with three-peak Gaussian fitting for 4Pi SRS, and single-peak Gaussian fitting for conventional SRS with FWHM results. (c, f, i) Lateral single-peak Gaussian fitted FWHM results for both x and y bead profiles. 80 nm PS beads were fitted with an average SNR of 7 (N=8). 200 nm PS beads were fitted with an average SNR of 15 (N=6). 500 nm PS beads were fitted with an average SNR of 30 (N=5). Scale bar = 1 µm.

For 200 nm beads, the axial profiles were object-limited, producing axial FWHMs consistent with bead size (Figure 2d-f). These results confirm object-limited axial resolutions of 236 ± 23 nm FWHM and lateral FWHM of 322 ± 43 nm and an SNR of 15. Finally, shown in Figure 2g-i, 500 nm beads produced object-limited axial and lateral resolutions of 500 nm. For objects >500 nm in diameter, the side lobes and central lobe overlap, resulting in a more complex profile. The sidelobes of these beads can be removed using deconvolution as shown in Supplemental Figure S4.

It is important to note that there is significant improvement in 4Pi-SRS signal over conventional SRS for small beads, much more than that observed in solution. This is because the interference increases the peak intensity by 2-fold, thus signal improves by 4-fold when beads are smaller or similar to the size of the 4Pi-PSF. For larger beads such as 500 nm beads, the signal increase is expected to be smaller. Nevertheless, our experiment still shows ∼3-fold signal increase.

### Image Lipid Droplet Distribution in Mammalian Cells

Lipid droplets (LDs) are central organelles that play essential roles in energy storage, metabolic regulation, and signaling of mammalian cells.^43,44^ In cancer cells, LD dynamics are tightly coupled to lipid synthesis, oxidative stress adaptation, and chemoresistance.^45,46^ Imaging lipid droplets with sub-diffraction resolution is essential for understanding their spatial organization and interaction with subcellular organelles. Although SRS has been widely used in studying lipid droplet distribution, composition, organization, and metabolic dynamics in many different cancer cells,^47–53^ conventional SRS is limited in axial resolution, hindering imaging of droplets <200 nm and accurate 3D volumetric mapping. Here, we apply 4Pi-SRS microscopy to image LD distribution in A549 cells, a non-small cell lung cancer cell line, taking advantage of its enhanced axial resolution and sensitivity to resolve the fine morphology and distribution of cytoplasmic lipid droplets.

Prior to quantitative analysis, the images were denoised using PureDenoise,^54^ suppressing photon shot noise while maintaining spectral and spatial details. The denoising did not alter the spectral or axial profiles as shown in Supplemental Figure S5. We first performed hyperspectral 4Pi-SRS imaging of A549 cells in the C–H region. By analyzing the spectrum of LDs, their high fatty acid content contributes to a strong signal at 2850 cm^-1^ (Fig. 3a-b). When comparing the maximum intensity projections from conventional SRS and 4Pi-SRS on the same intensity scale, we can observe significant contrast enhancement of LDs in 4Pi-SRS. This is largely because poorly resolved small LDs in conventional SRS have much higher signal to background ratio in 4Pi-SRS. Fig. 3c,i show the zoom-in of LDs and their lateral and axial profiles. We observed pronounced axial sharpening in the YZ planes and signal increase by ∼ 4-fold with 4Pi-SRS. For the small LD, Gaussian fit of cytoplasm background-subtracted axial profile yielded an axial FWHM of 1289 with conventional SRS versus 160 nm axial FWHM with 4Pi-SRS, while the lateral FWHM remained comparable at ∼294 nm (Fig. 3d,e). Deconvolution of the LD with the measured PSF of 142 nm suggests that the size of the LD is less than 100nm. A larger LD showed a similar trend (Fig. 3i,k), where conventional SRS resulted in an axial FWHM of 1450 nm due to cytoplasmic background, whereas 4Pi-SRS resolved the LD at 244 nm in addition to a ∼3-fold sensitivity improvement. Together, these examples illustrate a ∼7x improvement in axial resolution and up to 4-fold sensitivity improvement of 4Pi-SRS.

**Figure 3.**
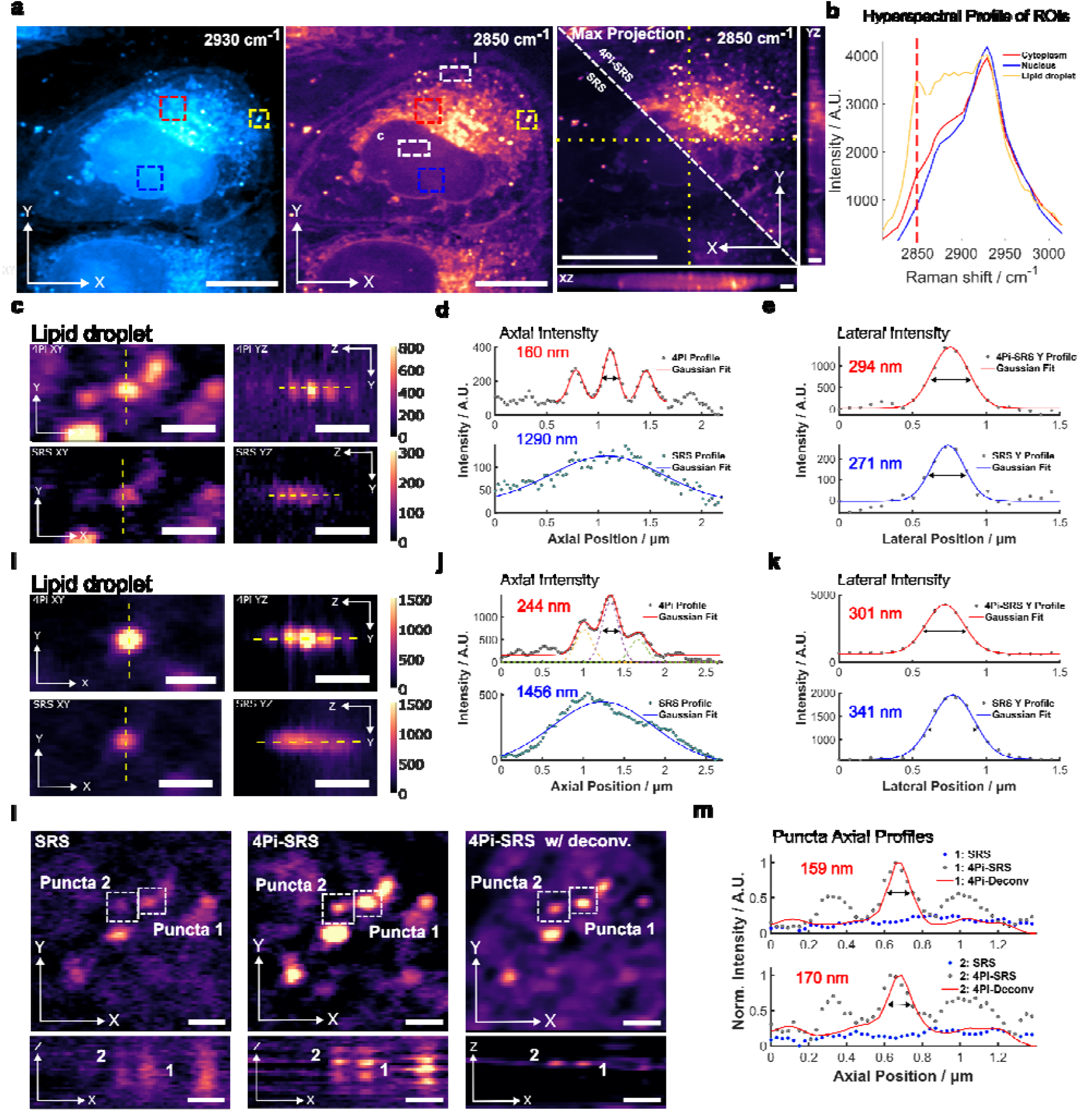
4Pi-SRS volumetric imaging of A549 cancer cell lipid droplet distribution. (a) SRS and 4Pi-SRS ima es acquired at 2930 and 2850 cm^-1^; maximum-intensity projections highlight improved contrast and structural detail with 4Pi-SRS. (b) Representative SRS spectra from cytoplasm and key organelles. (c,i) Zoomed-in regions (dashed boxes in a) shown in XY and YZ views. (d,j) Axial line profiles through LDs with background-subtracted Gaussian fits for conventional SRS and 4Pi-SRS. (e,k) Corresponding lateral profiles and Gaussian fitting. (l) XY and XZ comparisons among conventional SRS, 4Pi-SRS, and 4Pi-SRS after deconvolution. (m) Axial profiles illustrating side-lobe suppression with deconvolution. (a,b) XY, and XZ/YZ views scale bars = 10 µm and 2 µm, respectively. (c,i,l) Scale bar = 1 µm.

We next apply deconvolution to 4Pi-SRS to suppress side lobes and reduce axial ambiguity (Fig. 3l,m). Similar to 4Pi-fluorescence, axial interference introduces side lobes that degrade localization and quantification. Richardson-Lucy deconvolution is commonly used to remove side-lobes.^39,41,42^ We applied a non-blind hybrid L0-total variation deconvolution to suppress side lobes while preserving low-intensity chemical features.^40^ The experimental PSF was determined from the average of eight, 80 nm beads and parameters were chosen to balance suppression and artifact avoidance (Supplemental Figure S3). Under deconvolution, side-lobe reduction was observed, visualizing LD pairs localized within the cytoplasm between the coverslip and the nucleus with an axial FWHM of 159 nm and 170 nm for puncta 1 and 2, respectively. It is important to point out that deconvolution is sensitive to the choice of parameters and artifact can arise if parameters are not optimized carefully (Supplemental Figure S6). Some residual, low-intensity side-lobes remain. This is likely due to refractive-index heterogeneity in the cell, causing unbalanced sidelobes in 4Pi-SRS. Future work with a variable phase introduced to the PSF may further suppress the side lobes.

Overall, 4pi-SRS enables volumetric chemical imaging of subcellular lipid structures with ∼7-fold better axial resolution and 3-4 fold enhanced contrast, allowing the three-dimensional visualization of LDs that are otherwise unresolved by conventional SRS.

### Three-dimensional 4Pi-SRS imaging of *E. Coli* membranes

Next, we performed volumetric 4Pi-SRS imaging of *E. coli*, focusing on chemically distinct components of the cell envelope and extracellular fatty-acid bioproducts. The *E. coli* bacteria envelope comprises the inner membrane, a periplasm with peptidoglycan, and the outer membrane, with an overall thickness on the order of ∼60 nm. The diameter of *E. coli* is typically ∼0.5-0.7 μm, below the axial resolution of conventional SRS.^27,55,56^ These conditions pose a stringent test for the sensitivity and resolution of 4Pi-SRS imaging Prior to analyses, PureDenoise^54^ was used to suppress photon shot noise. Raw *E. coli* images and profiles can be found in Supplemental Figure S7. Hyperspectral 4Pi-SRS imaging across the CH window identified 2850 cm^-1^ as providing he best contrast for the cell envelope, whereas 2930 cm^-1^ emphasized cytoplasmic signal (Fig. 4a). In zoomed views (Fig. 4c,d,g,h), 4Pi-SRS improves the imaging contrast relative to SRS, allowing for much better contrast for the envelope, especially in the axial direction. Along cross-sections through the cell body, conventional SRS could not separate the two membranes; the axial profile is broadened with an FWHM of 1107 nm due to the diffraction limit. In contrast, 4Pi-SRS produced a multi-peak axial signature, arising from central and side lobes of each membrane. At the geometric center of the *E. coli*, the two sidelobes happen to overlap in the middle and give rise to a stronger peak. By fitting the axial profile with five Gaussian functions, we obtained a FWHM peak width for each membrane to be 220 nm. Applying deconvolution substantially suppressed the side lobes (Fig. 4e,f), cleanly resolving the two membranes in the short-axis cross-section with an axial center-to-center separation of ∼600 nm, underscoring the advantage of 4Pi-SRS for resolving biological nanostructures.

**Figure 4:**
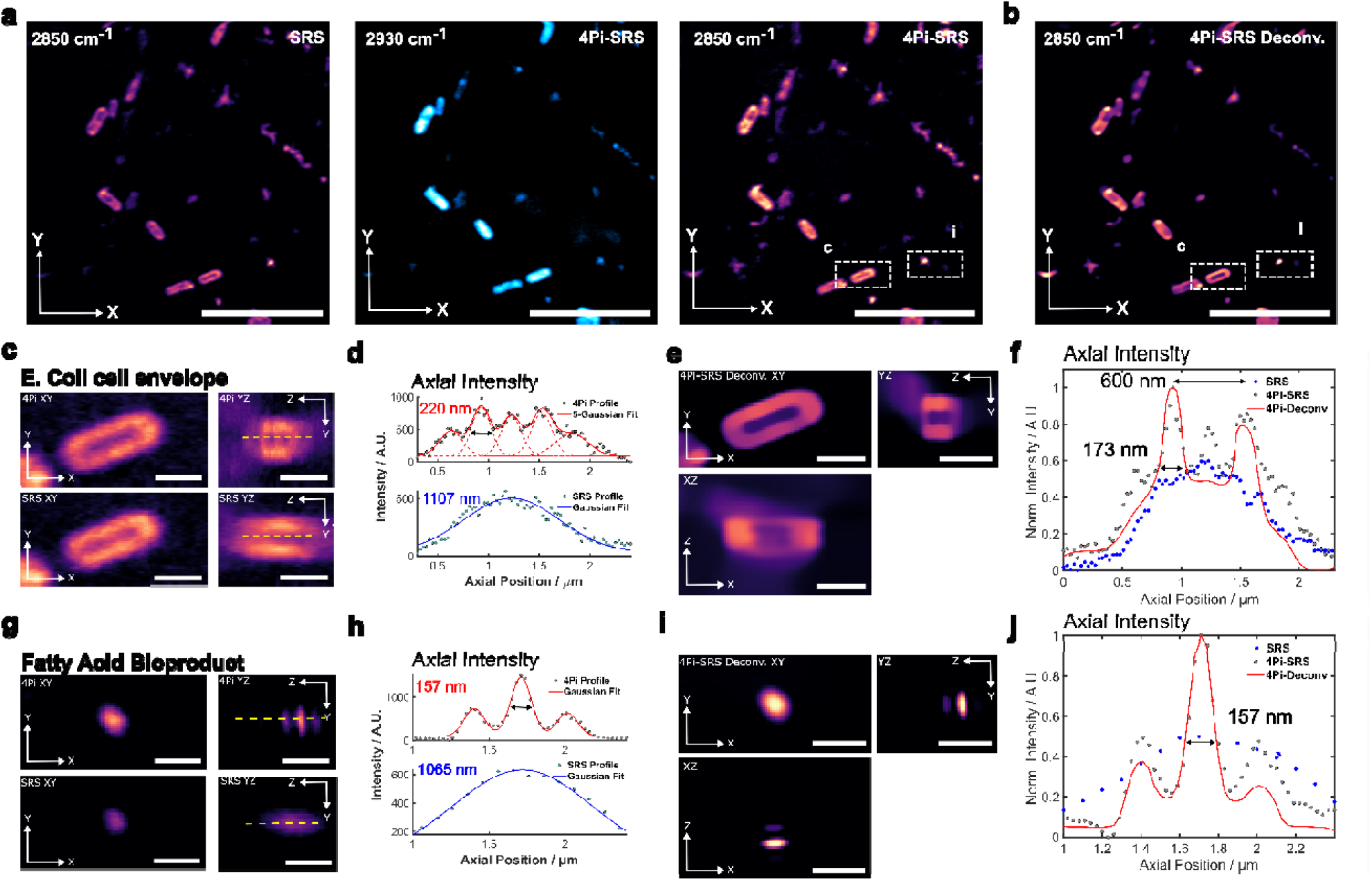
(a) Two-color 4Pi-SRS and 2850 cm^-1^ SRS images of *E. coli* with fatty acid bioproduct. (b) 2850 cm^-1^ 4Pi-SRS image after deconvolution. (a,b) scale bars = 10 µm. (c, i) Zoom-in of the dashed boxes showing XY and YZ planes for SRS and 4Pi-SRS. (d, j) Axial profiles of SRS and 4Pi-SRS line profiles. (e, k) 4Pi-SRS with deconvolution—XY, XZ, YZ planes of zoomed regions. (f, l) axial profiles of the 4Pi-SRS with deconvolution highlighting the *E. coli* cell membrane and fatty acid bioproduct. (c,e,g,i) scale bars = 1 µm.

We also resolved fatty acid bioproducts produced by *E. coli*. In conventional SRS, the axial profile was broad (FWHM 1065 nm) with low signal; 4Pi-SRS improved signal by ∼2× and super-resolved the feature to an axial FWHM of 157 nm (Fig. 4i–k). Deconvolution did not fully remove residual side lobes—likely due to phase mismatch between the deconvolution PSF and the sample’s interference phase—but the primary peak was sharply localized.

To probe the chemomorphological structure of the *E*. coli, we performed hyperspectral 4Pi-SRS analyses across the short axis of the *E. coli* (Supplemental Figure S8). Membrane regions displayed a dominant peak at 2850 cm□^1^, consistent with higher fatty acid content, whereas the cytoplasm exhibited a stronger 2930 cm□^1^ peak characteristic of proteins. In particular, ROIs 1 and 5 showed elevated 2850/2930 intensity ratios, highlighting lipid enrichment at the cell boundary.

Together, these data demonstrate chemical super-resolution of bacterial nanostructures with 4Pi-SRS, enabling separation of the *E. coli* membrane in three-dimensions along the short-axis and visualization of fatty-acid bioproducts at ∼157 nm axial FWHM.

## Discussion

Developing super-resolution chemical imaging is critical because many biological and biomedical questions depend on resolving nanoscale molecular organization without the use of labels. Conventional vibrational microscopy, including infrared and Raman imaging, though powerful in providing intrinsic molecular contrast, has been constrained by diffraction-limited resolution. This has restricted its ability to visualize subcellular nanostructures, such as lipid droplets, membrane domains, and metabolic assemblies, that play central roles in cellular physiology and disease. Addressing this limitation requires methods that can improve spatial resolution while preserving the chemical specificity and sensitivity. Super-resolution SRS is a promising approach, but unfortunately existing implementations mostly rely on the use of specialized probes, which negates the strength of label-free chemical imaging. Other label-free SRS approaches either have poor sensitivity, strong photodamage, or limited resolution improvement. Moreover, existing approaches focus on improving lateral resolution while the axial resolution was neglected, despite that it is the worst among the three dimensions.

In this work, we extend label-free SRS microscopy into the axial super-resolution regime by developing the 4Pi-SRS microscope. Across polymer beads, mammalian cells and bacteria, 4Pi-SRS sharpened the axial point-spread function and substantially increased image contrast, enabling visualization of chemical nanostructures that remain poorly resolved or substantially elongated in conventional SRS. We characterized the axial resolution of the 4Pi-SRS with 80 nm polystyrene beads to be ∼142 nm, the smallest among all demonstrated super-resolution SRS techniques. In A549 cells, lipid droplets that appeared axially broadened in regular SRS (FWHM >1 μm) were resolved by 4Pi-SRS to an axial width of 160 nm, suggesting that the size of the LDs are smaller than 100 nm. In *E. coli*, 4Pi-SRS separated cell envelope-associated features and visualized small fatty-acid bioproducts. Deconvolution was used to remove the sidelobes arising from the interference. Together, these results indicate an approximately seven-fold axial resolution improvement accompanied by up to 4-fold gains in sensitivity and contrast, allowing volumetric mapping of subcellular structures that were previously inaccessible.

Despite these advances, 4Pi-SRS still has practical constraints. Phase-stable alignment is required, and refractive-index heterogeneity can introduce phase distortions that generate residual side lobes after deconvolution. Sample thickness is limited by the short working distance between the two opposing high numerical aperture objectives, restricting applications primarily to monolayer cells or thin tissues. However, adaptive optics can restore phase fidelity in heterogeneous biological samples,^57^ and variable-phase deconvolution methods have been shown to suppress side lobes in 4Pi imaging.^39^ These approaches suggest potential pathways to overcoming the current technical barriers.

Possible future developments include integration with alternative resolution-enhancing methods such as visible excitation SRS and/or sample expansion to reach sub-100 nm lateral and axial resolution. Wavelength tuning to the CD signal region or fingerprint region will allow probing of biomolecules beyond lipids and proteins at resolutions far beyond current limits. In summary, 4Pi-SRS leverages the 4Pi interferometry to achieve robust axial super-resolution with enhanced sensitivity. With continued advances in phase control and resolution extension, this method offers a practical path toward three-dimensional chemical nanoscopy for mapping nanoscale structures and metabolic processes in various biological systems.

## Materials and Methods

### 4Pi-SRS Microscope system

We employed a ytterbium-doped femtosecond laser system (Flint-FL1, Light Conversion) to provide the Stokes output with a repetition rate of 75 MHz and a pulse duration of ∼80 femtoseconds. 80% of the laser output was frequency-doubled through a BBO crystal and used to pump an optical parametric oscillator (OPO) to generate a tunable pump output. For the polystyrene beads experiments, the pump wavelength was centered at 785 nm, and for imaging of A549 and *E. coli* cells, 793 nm. The Stokes beam was modulated by an electro-optical modulator (EO-AM-NR-C2, Thorlabs) at 17.5 MHz. Both pump and Stokes were stretched using grating stretcher systems to ∼2 picoseconds with matched chirp for an optimal spectral resolution of 14 cm-1. A motorized time delay stage (TRA25CC, Newport) on the pump output was used to implement hyperspectral scanning through spectral focusing. The pump and Stokes are spatially overlapped through a dichroic mirror (DMSP900T, Thorlabs). The two-color excitations were sent into the 4Pi-SRS system, 4f conjugated to the sample using a 2-axis scanning galvo mirror system, scan lens, and tube lens. For high-resolution imaging, we used matching high-NA water objectives (UPLSAPO60XW PSF; NA 1.2, Olympus). Nematic liquid crystal compensators were used for both the upper and lower arm chromatic dispersion compensation (LC1115-B, Thorlabs). The SRL signal was demodulated using a high-speed lock-in amplifier (Moku: Lab, Liquid Instruments). All imaging signals were acquired by a high-speed acquisition card (NI PXIe-6366, National Instruments).

### SRS and 4Pi-SRS Image Acquisition

For all experiments, we determined the laser power on the sample by measuring the power before the upper and lower objectives and multiplying it by the pre-calibrated objective transmission for both pump and Stokes wavelengths. The laser powers for the pump and Stokes were set to 20 mW and 20 mW, respectively, at the focus. For polystyrene, measurements were acquired with 8 μs pixel dwell time and 49 nm pixel size. For A549 cancer cells and *E. coli*, the measurements were acquired with 20 μs pixel dwell time, with 37.5 μm and 25 μm field of view, respectively. Volumetric images for A549 and *E*. coli were acquired with an axial spacing of 30 nm for biological experiments. For bead measurements of 80 nm, 200 nm, and 500 nm, the axial spacings used were 40 nm, 30 nm, and 110 nm, respectively. Hyperspectral imaging of A549 and *E. coli* cells was acquired with 40 hyperspectral frames, with 20 μs pixel dwell time covering the CH hyperspectral region (2800 cm^-1^ to 3000 cm^-1^).

### 4Pi-SRS PSF simulation

We simulated the 4Pi stimulated Raman scattering (SRS) point-spread function (PSF) using a vectorial diffraction framework, following Khonina et al.^37,38^ The simulation was carried out in MATLAB (GitHub link in code availability). The derivation for the 4Pi-SRS PSF simulation can be found in supplemental Note 1.

### 4Pi-SRS Denoising and Deconvolution

Prior to deconvolution, images were denoised using PureDenoise,^54^ to suppress photon shot noise while preserving spectral and structural features. The deconvolution code and example data were implemented in MATLAB (GitHub link in code availability) following Dong *et al*.^39,40^ The 4Pi-SRS PSF was experimentally measured from eight 80 nm beads. Bead volumes were denoised, cropped to equal dimensions, averaged, and normalized to unity sum. This bead-derived PSF was used as the input PSF to preserve the interference lobes critical for sidelobe removal in deconvolution. Reconstructions were initialized with Richardson-Lucy deconvolution^41,42^ to provide a physically consistent starting point, after which iterative maximum-a-posteriori (MAP) optimization with L0-TV regularization,^40^ was applied until the following convergence criteria:

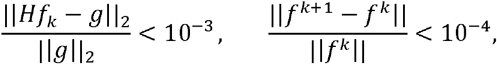

were satisfied, ensuring sidelobe suppression and artifact-free reconstructions.

### Preparation of polystyrene beads

Polystyrene beads with nominal diameters of 80 nm (64008, Polysciences), 200 nm (107, Phosphorex), and 500 nm (109, Phosphorex) were prepared. A 5 µL aliquot of bead suspension was mixed into 700 µL of 0.5% low-melting agarose (A4018, Sigma Aldrich) dissolved in DCO (151882, Sigma Aldrich). The mixtures were sandwiched between two cover glasses (CG15KH1, Thorlabs) and sealed with nail polish.

### Preparation of A549 cancer cells

A549 human lung carcinoma cells (A549 CCL-185, ATCC) were cultured in high-glucose DMEM (11-995-065, Fisher Scientific) supplemented with 10% fetal bovine serum (CH3007102, Fisher Scientific) and 1% penicillin–streptomycin (15-140-122, Fisher Scientific). Cells were maintained in a humidified incubator at 37 °C with 5% COC (*v/v*) and grown directly on 22 × 22 mm No. 1.5H coverslips (CG15CH, Thorlabs). After 48 h, cells were fixed in 4% paraformaldehyde (AAJ19943K2, Fisher Scientific) for 20 min and washed three times with phosphate-buffered saline (10010023, Thermo Fisher). For imaging, each coverslip was sandwiched with a second coverslip (CG15KH1, Thorlabs) and sealed with nail polish.

### Preparation of *E. Coli* cells

*E. coli* K-12 MG1655 glycerol stock was streaked on LB agar plates without antibiotics and incubated overnight at 30 °C until visible colonies formed. Sterile culture tubes containing 5 mL of LB medium were prepared, and individual colonies were used to inoculate the tubes. Cultures were grown overnight (12–16 h) at 37 °C in an orbital shaker incubator.

One milliliter of each culture was transferred to a 1.7 mL microcentrifuge tube (T3-300-500-Y, Stellar Scientific) and centrifuged at 10,000 × g for 1 min. The supernatant was discarded, and the pellet was washed three times with 1 mL phosphate-buffered saline (PBS; 10010023, Thermo Fisher). For fixation, the bacterial pellet was resuspended in 1 mL of 4% paraformaldehyde (AAJ19943K2, Fisher Scientific) and incubated on ice for 10 min. The fixed *E. coli* were then washed three times with PBS (10010023, Thermo Fisher).

## ACKNOWLEDGEMENTS

This work was financially supported by NSF CAREER 1846503 to D. F. The SRS microscope was developed with partial support from NIH R35GM133435 to DF. We thank the University of Washington, Department of Chemistry, machine shop, for the equipment to develop the custom-machined 4Pi-SRS microscope elements.

## Author information

## Contributions

D.F. conceived the project. J.K., Z.E., and D.F., constructed the 4Pi-SRS system. D.F. and J.K. developed the 4Pi-SRS simulation modeling. E.D. prepared A549 cancer cells and J.K. performed 4Pi-SRS experiments. N.M. prepared *E. Coli* and J.K. performed 4Pi-SRS experiments. J.K. performed denoising and deconvolution analysis. J.K. and D.F. wrote the manuscript with input and approval from all authors. D.F. supervised the project.

