## supplemental figures for "4-Pi Stimulated Raman Scattering for Label-free Super-resolution Chemical Imaging"

### Table of Contents

Supplemental Notes

1. 4Pi-SRS PSF simulation…………………………………….…………………………….....2

2. Deconvolution pipeline……....…………………………………………………………….…...3

3. 4Pi-SRS hyperspectral analysis of *E. coli* membrane content heterogeneity……………..……7

Supplemental Figures

Figure S1. Deconvolution pipeline………………..………..………………..……...…….…...3

Figure S2. Limit of detection (LOD) for 4Pi-SRS…………………………….…………………..4

Figure S3. 4Pi-SRS images of 80 nm beads…………………..………………..…….….……4

Figure S4. Deconvolution of beads……………………………………………………..……...5

Figure S5. Effects of denoising on hyperspectral and lipid droplet profiles……………………....5

Figure S6. Characterization of deconvolution artifacts…………….…………..…..…………6

Figure S7. Raw images of *E. coli* cell envelope and fatty acid bioproduct…………………..……6

Figure S8. 4Pi-SRS hyperspectral analysis of *E. coli* membrane content heterogeneity…......…...7

**Supplementary notes 1**: **4Pi-SRS PSF simulation.** We simulated the 4Pi stimulated Raman scattering (SRS) point-spread function (PSF) using a vectorial diffraction framework, following *Khonina et al.*[1,2] The simulation was carried out in MATLAB (GitHub link in code availability). The pump and Stokes beams were assumed to be x-linearly polarized, simplifying the simulation while retaining generality. The objective numerical aperture (NA) was set to 1.2 with an immersion refractive index of n = 1.33, yielding a maximum convergence angle:

$$\alpha = arcsin(\frac{NA}{n})$$

The pump and Stokes wavelengths were chosen as $\lambda_{pump}$ = 793 nm, and $\lambda_{Stokes}$ = 1030 nm, with corresponding wavenumbers

$$k_{pump} = \frac{2\pi n}{\lambda_{pump}}, \qquad k_{Stokes} = \frac{2\pi n}{\lambda_{Stokes}}$$

The vectorial fields were calculated using Debye—Wolf angular spectrum approach. The pupil was parameterized in polar coordinates (θ, $\varphi$), with $\theta \epsilon [0, \alpha]$ and $\varphi \epsilon [0,2\pi]$. The pupil was uniformly sampled with with **Nθ** = 50, and **Nφ** = 50, yielding angular steps $\Delta\theta$ and $\Delta\varphi$. Given the x-linear polarization input beam

$$P = \begin{bmatrix} 1 \\ 0 \\ 0 \end{bmatrix}$$

The field after high-NA focusing was obtained using the Debye—Wolf polarization mapping:

$$P'(\theta, \varphi) = V(\theta, \varphi)P$$

where $V(\theta, \phi)$ is the transmission function of the incoming beam. The polarization matrix reduces to:

$$P'(\theta, \varphi) = \begin{bmatrix} 1 + \cos^2 \varphi \, (\cos\theta - 1) \\ \sin\varphi \cos\varphi \, (\cos\theta - 1) \\ - \cos\varphi \sin\theta \end{bmatrix}$$

An apodization factor $sin\theta\sqrt{cos\theta}$ was applied to ensure aplanatic weighting, corresponding to a system free of spherical and coma aberrations.

With this, the electric field in the lower $(+z)$ and upper $(-z)$ arms were computed via angular spectrum integration:

$$E^{(+)}(\theta, \varphi) = \int_0^{2\pi} \int_0^{\alpha} sin\theta\sqrt{\cos\theta}\, e^{ik[zcos\theta + x\sin\theta\cos\varphi + y\sin\theta\sin\varphi]} \, d\theta \, d\varphi$$

$$E^{(-)}(\theta, \varphi) = \int_0^{2\pi} \int_0^{\alpha} sin\theta\sqrt{\cos\theta}\, e^{ik[-zcos\theta + x\sin\theta\cos\varphi + y\sin\theta\sin\varphi]} \, d\theta \, d\varphi$$

The total fields for pump and Stokes were obtained by coherently adding the lower and upper arm contributions.[3,4] The 4Pi-SRS excitation PSF was then expressed as:

$$SRS_{4Pi} \propto \sigma |E^{+}_{pump} + E^{-}_{pump}|^2 * |E^{+}_{Stokes} + E^{-}_{Stokes}|^2$$

where $\sigma$ denotes the Raman scattering cross-section.

**Supplementary Notes 2: Deconvolution pipeline.** Supplemental Figure 4 illustrates the deconvolution pipeline starting from a raw 4Pi-SRS volume, denoising, and deconvolution of the image volume. Deconvolution pairs well with 4Pi interferometry, as axial side lobes present in the axial PSF lead to axial ambiguities. Deconvolution helps suppress or remove the side lobes fully, removing the associated axial ambiguities. PureDenoise, with 4 spin cycles and 3 multiframe, was used to improve the image SNR before deconvolution. The denoised average PSF was used to represent the experimental PSF for the microscope. An initial 5 iterations of Richardson-Lucy (RL) deconvolution,[5,6] are applied to warm start the deconvolution process. We applied non-blind hybrid L0-TV deconvolution,[7] with the inputs being the denoised image and the experimental PSF. For the deconvolution, if the data residual of the denoised image and the deconvolution iteration reaches below the threshold of $10^{-3}$ or for the relative change between iterations being less than $10^{-4}$, then the deconvolution has reached the convergence tolerance and is terminated.

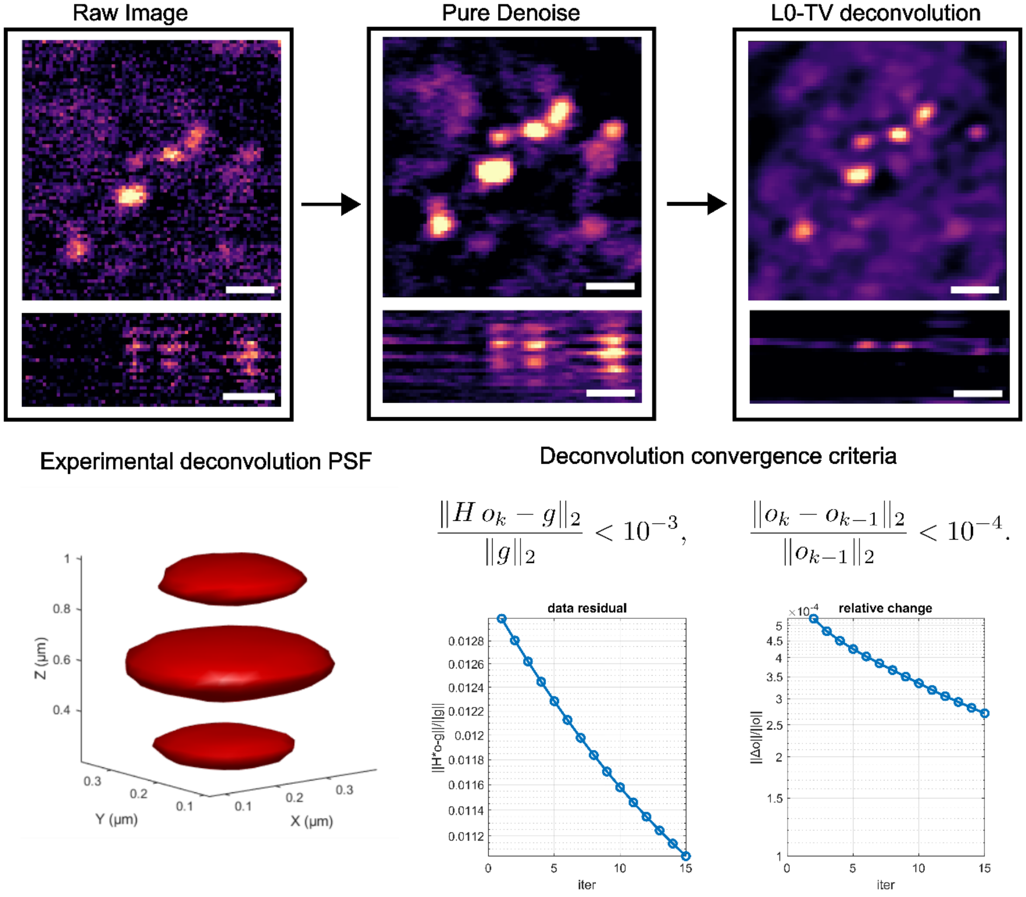


**Figure S1: Deconvolution pipeline.** (a) Flowchart of the processing sequence. The raw image is first denoised using PureDenoise, followed by non-blind hybrid $L_0$–TV deconvolution with the denoised volume and the experimental 4Pi-SRS PSF. Deconvolution terminates when the data residual is $< 10^{-3}$ and the relative change is $< 10^{-4}$, indicating convergence.

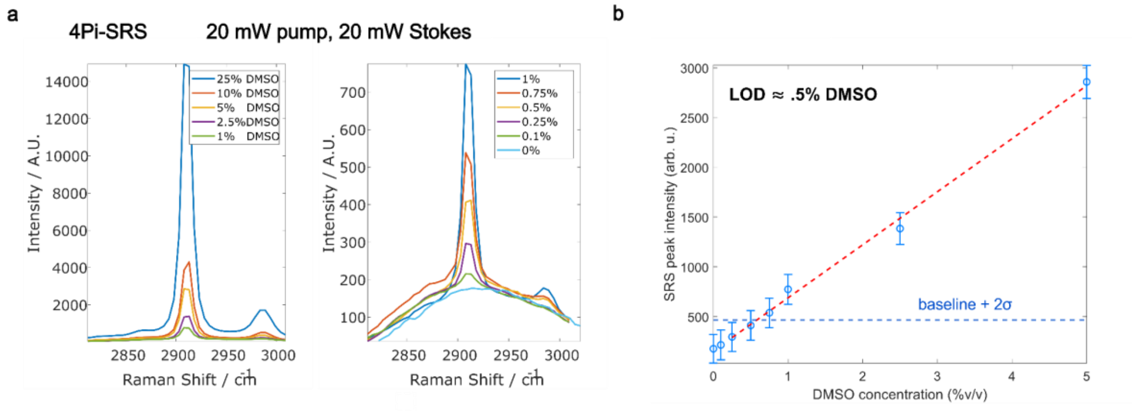


**Figure S2: Limit of detection (LOD) for 4Pi-SRS:** (a) 4Pi-SRS spectra of DMSO serial dilution in water. A separate zoomed view of the lower DMSO concentration spectra. (b) LOD was calculated from the maximum intensity of 0% DMSO + 2σ. σ is defined as the standard deviation of the bounding area where the mean intensity was observed (n = 400 pixels). Pixel dwell times of 4 µs were used during sensitivity measurement.

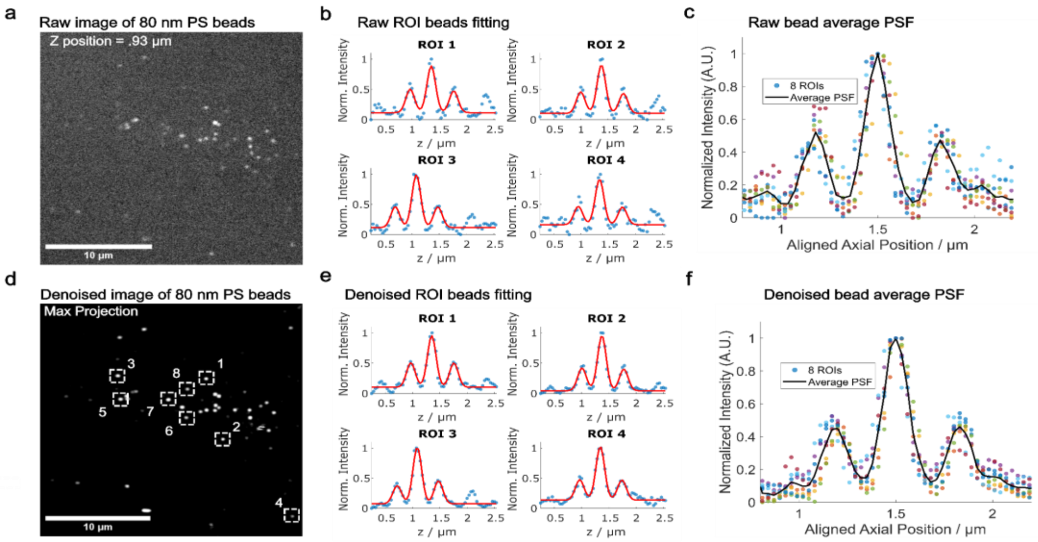


**Figure S3: 4Pi-SRS images of 80 nm beads:** (a) raw volumetric 4Pi-SRS image of 80 nm polystyrene (PS) beads taken at the z-slice = .93 µm. (b) 4 axial profiles of raw 80 nm beads. (c) The average raw 4Pi-SFS PSF of 8, 80 nm beads. (d) Denoised 4Pi-SRS max-projection image showing the location of beads 1-8 taken for axial profile fitting. (e) 4 axial profiles of denoised 80 nm beads. (f) Denoised average 4Pi-SRS PSF of 8, 80 nm beads.

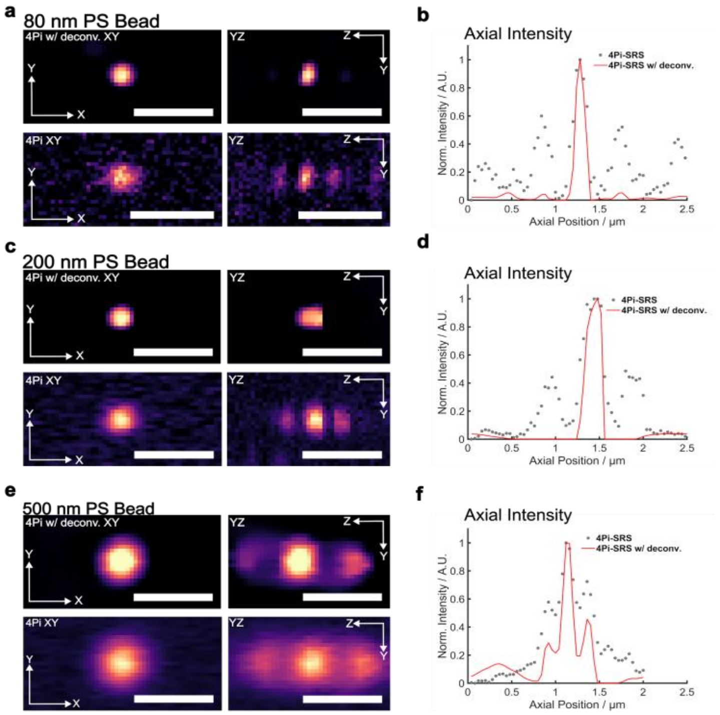


**Figure S4: Deconvolution of beads.** (a,c,e) Comparison of 4Pi and 4Pi-SRS with deconvolution XY and YZ profiles. (b, e, h) Overlaid axial profiles of 4Pi-SRS and 4Pi-SRS with deconvolution.

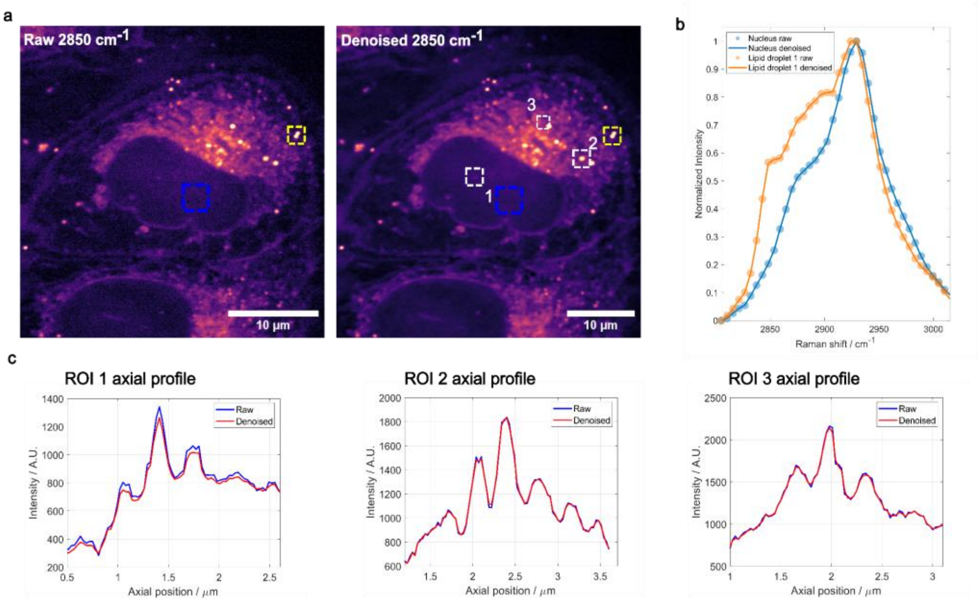


**Figure S5: Effects of denoising on A549 hyperspectral and lipid droplet profiles.** (a) Raw 4Pi-SRS image and a denoised max projection of an A549 cancer cell probing 2850 cm$^{-1}$. (b) overlaid raw and denoised hyperspectral line profiles of key organelle structures. (c) Axial line profiles of raw and denoised lipid droplets. The raw and denoised profiles show the same features, validating that the denoising is not affecting the spectra or axial profiles for key organelles or lipid droplets. PureDenoise parameters were set to 4 cycle-spins and 3 multiframe for denoising.

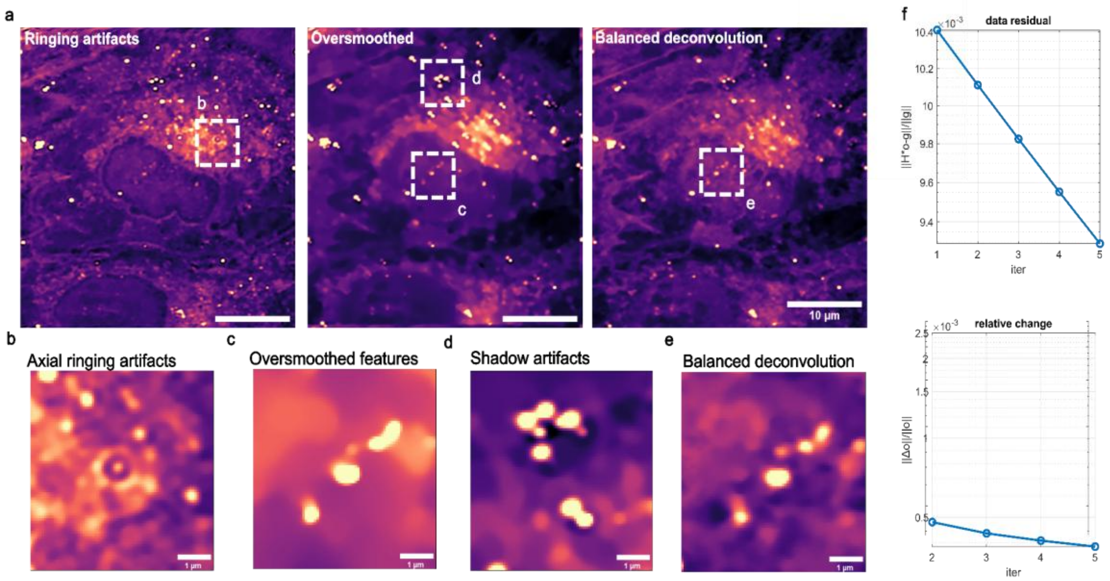


**Figure S6: Characterization of deconvolution artifacts.** (a) Deconvolved A549 cell images showing examples of imaging artifacts alongside a balanced deconvolution result. (b) Zoomed-in XY view illustrating axial ringing artifacts caused by excessive Richardson–Lucy (RL) iterations. (c) Excessive total variation (TV) regularization weight oversmooths low-intensity features. (d) Too many $L_0$–TV iterations can produce shadow artifacts adjacent to high-intensity structures, such as lipid droplets. (e) Balanced deconvolution preserves low-intensity features, avoids shadow artifacts, and suppresses axial side lobes. (f) Deconvolution residuals and relative change are tracked at each iteration to determine the termination point.

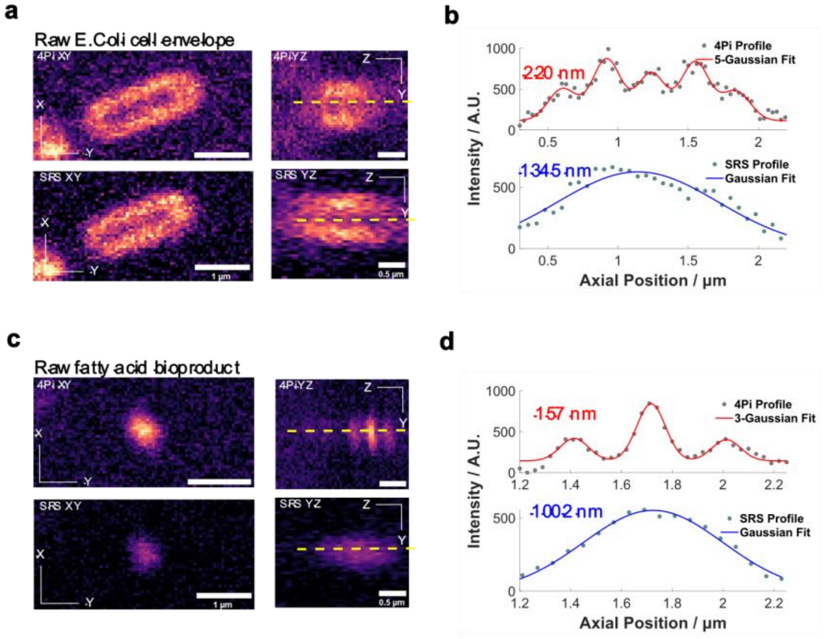


**Figure S7: Raw images of *E. coli* and fatty acid bioproduct.**(a) XY and YZ SRS and 4Pi-SRS views at 2850 $cm^{-1}$ of the *E. coli* cell envelope. (b) Axial profiles of the *E. coli* cell envelope, with 4Pi-SRS data fitted using a 5-peak Gaussian model and SRS data fitted using a single Gaussian peak. (c) XY and YZ SRS and 4Pi-SRS views at 2850 $cm^{-1}$ of the *E. coli* fatty acid bioproduct. (d) Axial profiles of the fatty acid bioproduct, with 4Pi-SRS data fitted using a 3-peak Gaussian model and SRS data fitted using a single Gaussian peak.

**Supplementary notes 3: 4Pi-SRS hyperspectral analysis of *E. coli* membrane content heterogeneity.** Figure S8 presents the spectrum variation along the short axis of *E. coli*. The membrane region exhibits a strong peak at 2850 cm⁻¹, indicative of higher fatty acid content, whereas the cytoplasm shows a pronounced 2930 cm⁻¹ peak associated with protein content. In ROIs 1 and 5, the spectra display a high 2850 cm⁻¹-to-2930 cm⁻¹ intensity ratio originating from the cell membrane region due to increased fatty acid content at the membrane.[9]

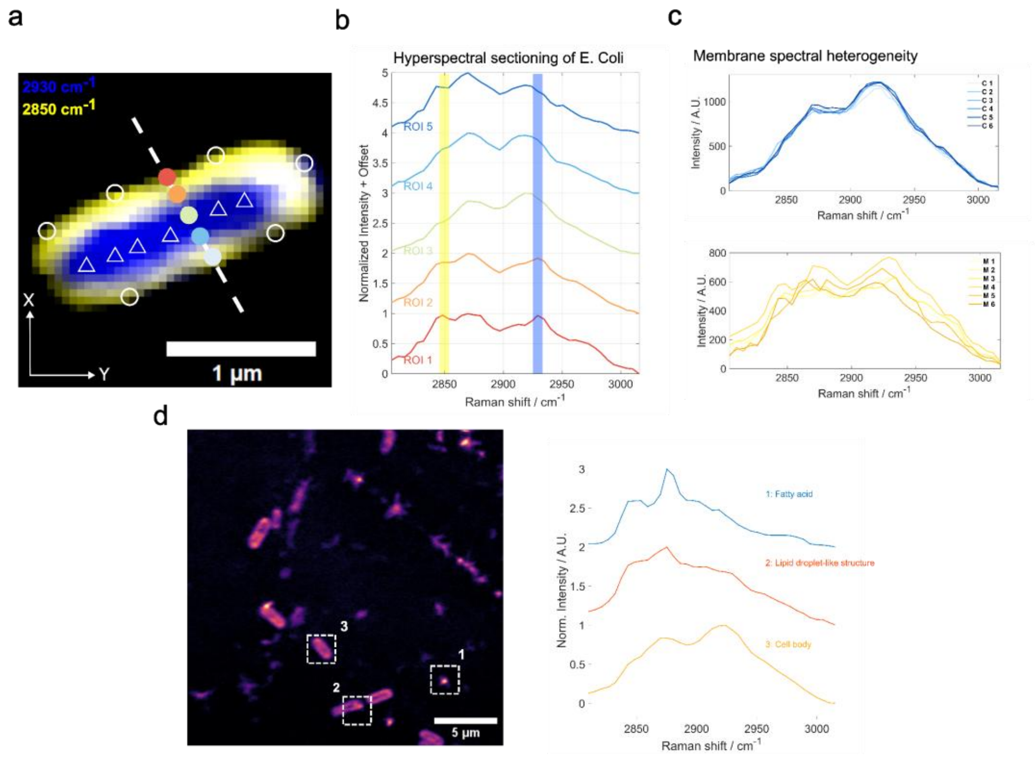


**Figure S8: 4Pi-SRS hyperspectral analysis of *E. coli* membrane content heterogeneity.** (a) Two-color image of an *E. coli* cell, where yellow indicates regions with dominant 2850 cm⁻¹ and 2930 cm⁻¹ signals. The white line and dots mark the hyperspectral sectioning path. Circles indicate positions where membrane spectral profiles were acquired; triangles indicate cytoplasmic sampling points. (b) Normalized profiles of the CH region (2800 cm$^{-1}$ – 3000 cm$^{-1}$) spectral profiles along the cross-section. (c) Spectral line profiles of the membrane (circles) and cytoplasm (triangles), corresponding to the positions marked in panel (a). (d) Hyperspectral image highlighting spectral features associated with fatty acid bioproducts, revealing lipid droplet–like structures within the *E. coli* cell body.
